# Telomere Orchestrates Encapsulation of Mitotic Chromosomes

**DOI:** 10.1101/2025.04.13.648587

**Authors:** PoAn Brian Yang, Takashi Mikawa

## Abstract

Telomere is crucial for continued cell proliferation and survival. Using high-temporal microscopy, we show that in mitosis it provides an initiation site for reassembly of barrier/scaffolding proteins during nuclear envelop reassembly. Subsequent spread of these proteins onto decondensing chromosomes completes encapsulation of the entire chromosome set. Depletion of telomere-tip components, TRF1 and TRF2, randomizes nuclear envelope reassembly initiation sites leading to multiple micronuclei production. The data show that telomere tip orchestrates this precisely timed program of nuclear envelop re-assembly.

**One Sentence Summary:** Yang and Mikawa investigated the role of telomere in initiating and forming a single nucleus.

## Main Text

In majority of higher eukaryotes, cell division involves open mitosis, which requires the complete breakdown and subsequent reassembly of the nuclear compartment (1) to encapsulate replicated chromosome set into each daughter nucleus (2). This evolutionarily conserved cell division program secures faithful delivery of genetic information to daughter cells. Incomplete encapsulation of the chromosome set leads to micronuclei, which contributes to massive chromosome recombination and damage known as chromothripsis (3). Despite its significance, how the reassembly process is precisely programed remains controversial in part due to complex multi-step assembly within a short time frame (4, 5). The outer layer of the nucleus, the nuclear envelope (NE), is known to be involved in maintenance (6) and regulation (7) of telomeres, which begs the question of whether the association of the two plays a role during the reformation of NE following cell division. Interestingly, although telomeres are localized mostly in the interior 50% of the human cell nucleus during interphase (8), they are located peripherally near the NE following mitosis (9) before relocating to the interior of the nucleus. Studies were done in other models, and during all phases of the cell cycle in budding yeast and mouse cells, telomeres on each chromosome not only resides in the peripheral regions of the nucleus, they are tethered to the NE (8, 10). Despite these correlations, the potential role of telomeres in reassembly of the nucleus during mitosis has largely remain untouched.

We began by investigating the location of various NE associated proteins throughout the reassembly of the nuclear compartment by analyzing high resolution live images. Nuclear compartment reassembly begins at early telophase through dephosphorylation of NE proteins (11), therefore, we focused on capturing live images during anaphase and telophase. As reported, ER and many NE resident proteins are excluded from the core region of the chromosome mass due to the dense mitotic spindles during anaphase and telophase (12, 13). However, by striking contrast, proteins such as lamin and barrier to autointegration factor (BAF) are present throughout the cell cytoplasm (Fig. 1A). Using high resolution live-imaging, we collected movies of mitotic cells with different NE components tagged with fluorescent labels and tracked their DNA volume using live-imaging dye SiR-DNA (14). Using Imaris (BitPlane) to reconstruct each nuclei in 3D (Fig 1B, movie S1), we measured the volume change normalized to each nucleus’ minimum volume. Utilizing chromosome condensation levels as a reference, we then compared relative location and timing of NE protein reassembly on chromosomes during mitosis.

**Fig. 1.**
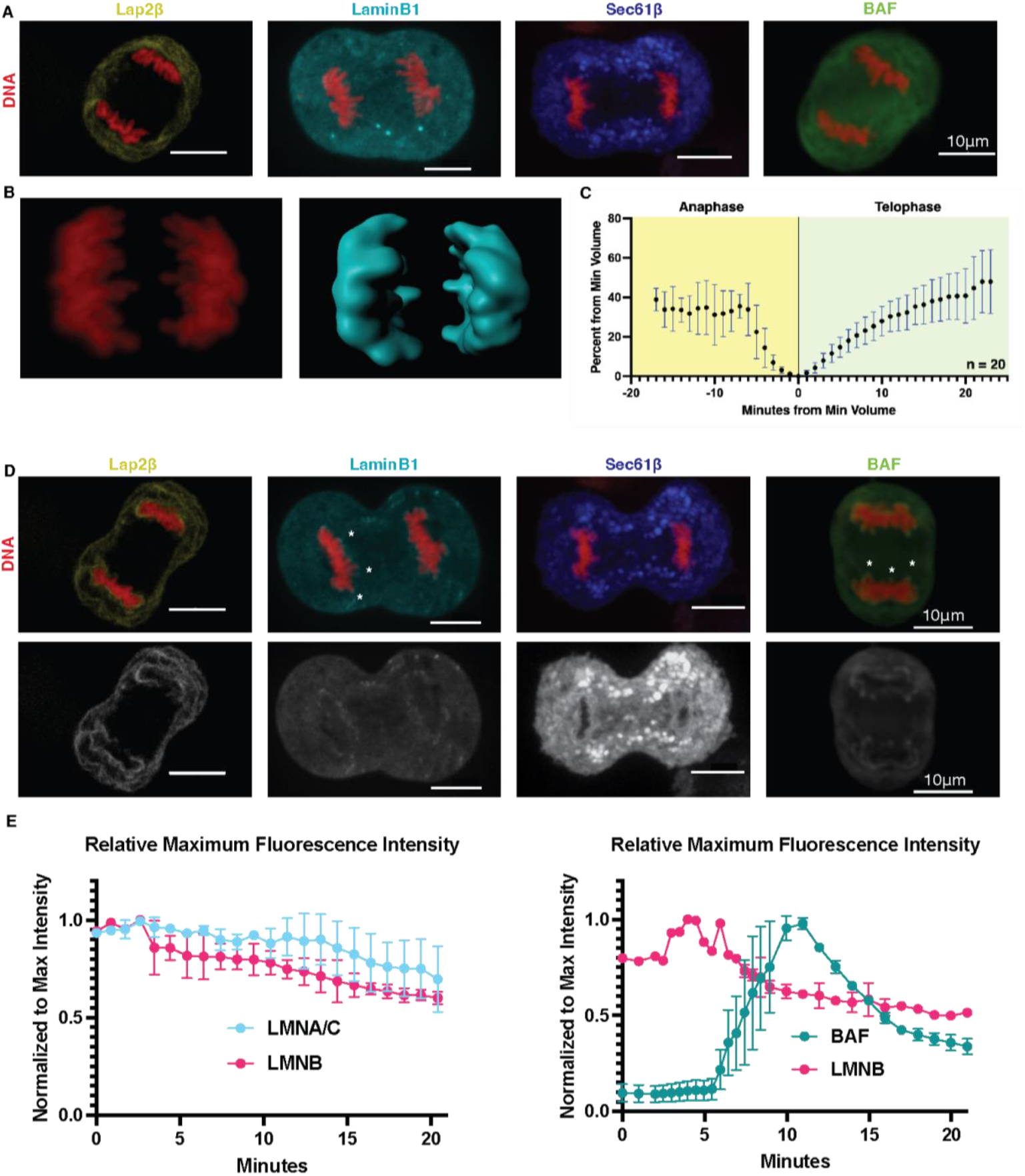
Comparison of NE Proteins’ Interaction with Chromosomes During NE Reformation. (**A**) Optical section of mitotic cells expressing different NE protein markers (Lap2β-GFP shown in yellow, LaminB1-RFP-T shown in cyan, Sec61β-GFP shown in blue, or BAF-GFP shown in green) are compared in late anaphase. Scale bar = 10um. (**B**) Example of DNA volume calculation using Imaris. DNA in red is used to generate a surface in cyan using the same threshold for all samples. (**C**) Relative DNA volume level during cell division was measured for 20 nuclei, and combined result shows distinguishable pattern of volume change from anaphase to telophase. Black dots represent mean, error bars in blue. (**D**) Optical section of mitotic cells expressing different NE protein markers are compared in early telophase. Accumulation of LMNB1 and BAF at the tip of chromosome arms can be observed, and indicated by asterisks (*). Scale bar = 10 um. (**E**) Maximum Fluorescence Intensity was measured in cells co-expressing LMNB1 and LMNA/C or BAF, after normalizing the highest signal to 1, comparison between the three shows that LMNB1 and LMNA/C reaches highest local density before BAF.

Consistent with previously reported studies (15), although DNA condensation begin once cell cycle reaches mitosis, maximum DNA compaction occurs at the end of anaphase, when reassembly of NE components began, and maximum condensation was then followed by de-condensation that ushers in telophase (Fig 1C). Our live imaging of major NE components during this critical time window for NE reformation revealed distinct reassembly patterns for several proteins in late anaphase. Most noticeably, LMNB and BAF, both of which are distributed throughout the cytoplasm during metaphase and early anaphase, appears to concentrate at the tip of chromosome arms during late anaphase (Fig 1D, movie S2, S3, and S4) before encapsulating the entire chromosome set. Since the site of accumulation could have a major impact on the formation of a single nucleus due to barrier function of LMNB and BAF (16, 17), we compared relative expression levels of Lamin and BAF. Comparing the LMNA/C, LMNB, and BAF by co-expressing pairs in the same cell, we see that both LMNA and LMNB reaches maximum local concentration before BAF, although the order of LMNA and LMNB can’t be confidently determined based on this method (Fig 1E, S1).

To verify that this selective localization of Lamin protein was not cell specific, we imaged multiple cell types expressing fluorescently labeled LMNB1, and localization was consistently seen at the tip before spreading (Fig B1). The localization of nuclear lamina to the chromosome tip can be caused by 1) suppression of these protein’s access to the inter-chromosome space, 2) attraction of barrier protein at the telomere tips, or 3) both. We began by testing the first possibility of telomere localization being due to the lack of accessibility to the inter-chromosomal space. We utilized taxol with reversine, stabilizing microtubules that keeps individual chromosomes apart and bypassing the spindle assembly checkpoint, thereby allowing encapsulation of individual chromosomes in treated cells (movie S5). We then visualized encapsulation pattern of LMNB on individual chromosomes, and similar to untreated cells, LMNB1 first localized at the telomere region, then spread toward the centromere region labeled by CENP-A (Fig 2B), despite the chromosomes being separated. Since BAF is thought to play a crucial role in chromosome zipping and thus inhibiting access of NE to inter-chromosomal space (17), we also depleted BAF using siBAF at 100nM concentration (Fig S2A). We first observed that de-condensation of mitotic chromosome of resulting BAF-depleted cells were greatly hampered (Fig S2B, C, and movie S6), yet with the depletion of BAF, LMNB1 still localized to the chromosome tip (Fig S2D and movie S7), demonstrating that DNA zipping was not responsible for telomere association of LMNB1. These results demonstrate that LMNB1 can spread to other regions of the chromosome and yet displays preferential binding initiation sites at telomere tips.

**Fig. 2.**
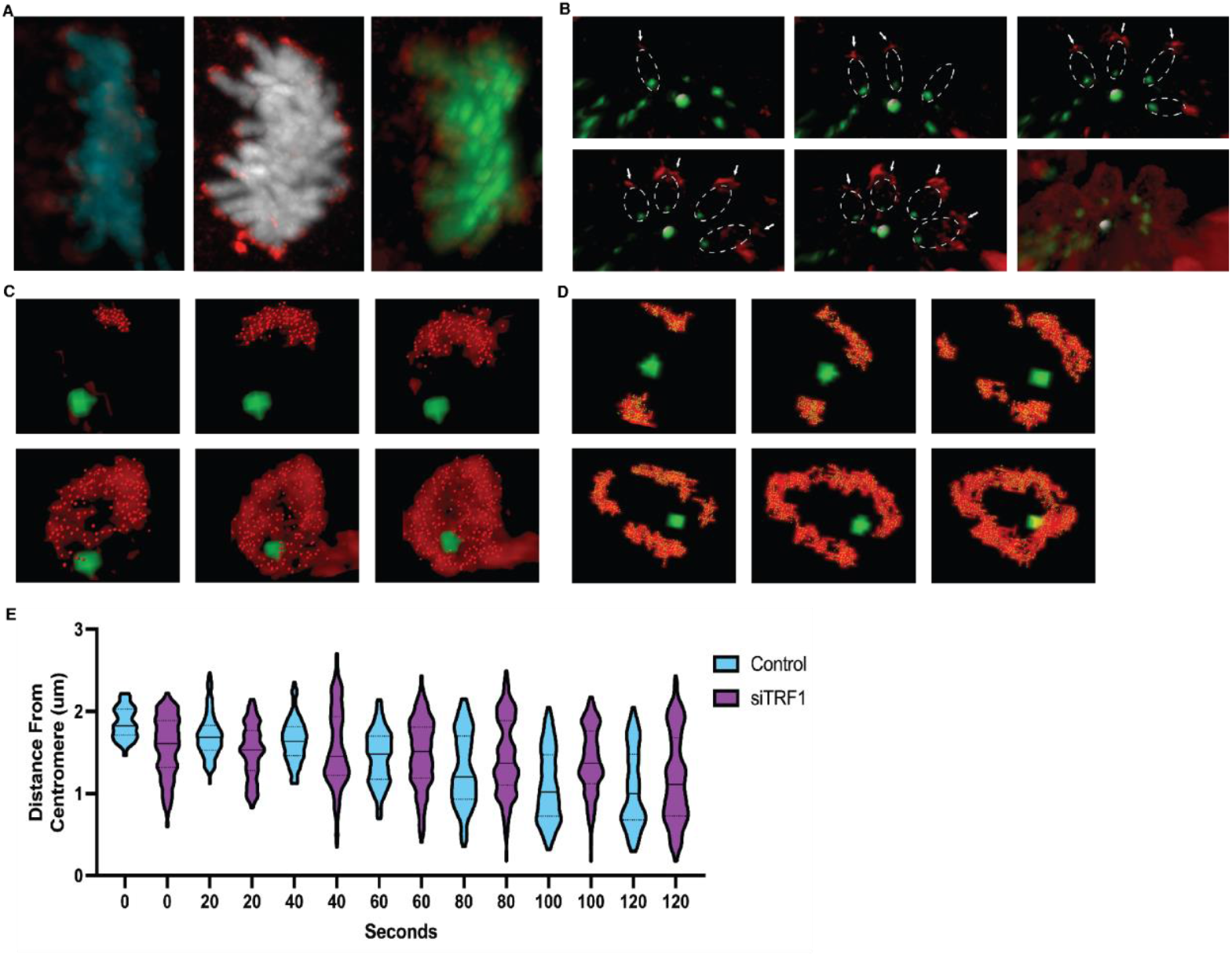
Accumulation of lamin at chromosome tip is observed in multiple cell types, but tip accumulation is disrupted by TRF1 inhibition. (**A**) LMNB1 (red) is seen to accumulate at the telomere region in different cell types: U2OS (Cyan) with GFP tagged endogenously to LMNB1, MCF10AT (Gray) with exogenous expression of LMNB1-RFP-T, and RPE-1 (Green) with exogenous expression of LMNB1-RFP-T. RPE-1 cells also express CENPA-GFP, which marks the centromere, and LMNB1 is seen to accumulate distal to the centromere. (**B**) Individual chromosomes around centromere (white) are encapsulated by lamin as it spreads from telomere toward centromere region. (**C**) Snapshots from live imaging of a control RPE-L chromosome with taxol and reversine undergoing mitosis. Going from top left to right. (**D**) Snapshots from live imaging of siTRF1 RPE-L chromosome with taxol and reversine undergoing mitosis. (**E**) Area of LMNB1 (red) from (**C**) and (**D**) are randomly seeded with points in each frame (n>1000), and distance of each point from centromere (green) is calculated. (**E**) Violin plot of control vs siTRF1 treated cells showing distance of lamin from centromere in each frame (n>5000). Width represents frequency of points at each distance.

Our results are consistent with the model wherein rather than inhibitory signals at the chromosome arm, attractive force at the chromosome tip patterns an initiation site for LMNB1 reassembly. Human lamins are attached to the nuclear matrix through shelterin interaction (18), and Telomere-Association-Assay showed more specifically that LMNB1 co-precipitated with shelterin component TRF1 (19). TRF1 has also been shown to bind to telomeric double-stranded 5’-TTAGGG-3’ repeat (20) and recruits other shelterin components to the telomere region (21). We therefore tested a potential involvement of the shelterin complex in recruitment of LMNB1 at telomere tips at the initiation of NE reassembly. We targeted TRF1 using siRNA knockdown (Fig S3A). Compared to control, cells with TRF1 knockdown displayed signs of telomere fusion (Fig S3B and movie S8) as previously reported (22). Furthermore, when treated with taxol and reversine, compared to control (Fig 2C) individual chromosomes in TRF1 knockdown group now displayed randomized LMNB1 localization and spread (Fig 2D and E). These results support the idea that the telomere plays a role in patterning initiation sites for recruiting nuclear lamina to the chromosome.

To assess the impact of NE protein mis-localization, we took large fields of image after inhibiting TRF1, TRF2, TRF1 and 2, or BAF using RNAi. We then analyzed the images (Data S1) and quantified the change in nuclear form using ImageJ with the FUJI plugin (23), and instances of micronuclei formation using CellProfiler 3.0 (24). The analysis revealed a wide range of phenotypes from shelterin complex disruption (Fig 3A, B, C, and D), with BAF having the biggest effect on nuclear circularity or nuclear form (Fig 3E), while the knockdown of TRF1, also double knockdown of TRF1 and TRF2 resulting in the highest instance of micronuclei formation (Fig 3F). Live images of individual siTRF1 treated cells undergoing mitosis (Fig 3G and movie S9) revealed that multinucleation occurs in cells exhibiting telomere fusion, a phenotype associated with shelterin disruption (25). These data suggest that shelterin complex depletion induced random LMNB1 localization prior to de-condensation gives rise to precocious ensheathing of individual chromosomes resulting in the formation of micronuclei. Although TRF1 knockdown could have a cascade of effects, including the loss of D-loop structure (26), telomere sequence recombination (27), and de-localization of multiple proteins at the telomere region (28), nevertheless, our work demonstrates how disruption of the telomere tip integrity has a direct and noticeable impact on the secure encapsulation of the entire chromosome set into a single nucleus. Recently, degradation of TRF1/2 through nucleotide-based proteolysis-targeting chimera (PROTEC) is shown to be effective in suppressing tumor cell proliferation, but has mild effect in normal cells from proliferation assays (29), and our study suggest that cancer cells may be more susceptible to TRF1/2 degradation due to frequent cell divisions that can lead to multinucleation.

**Fig. 3.**
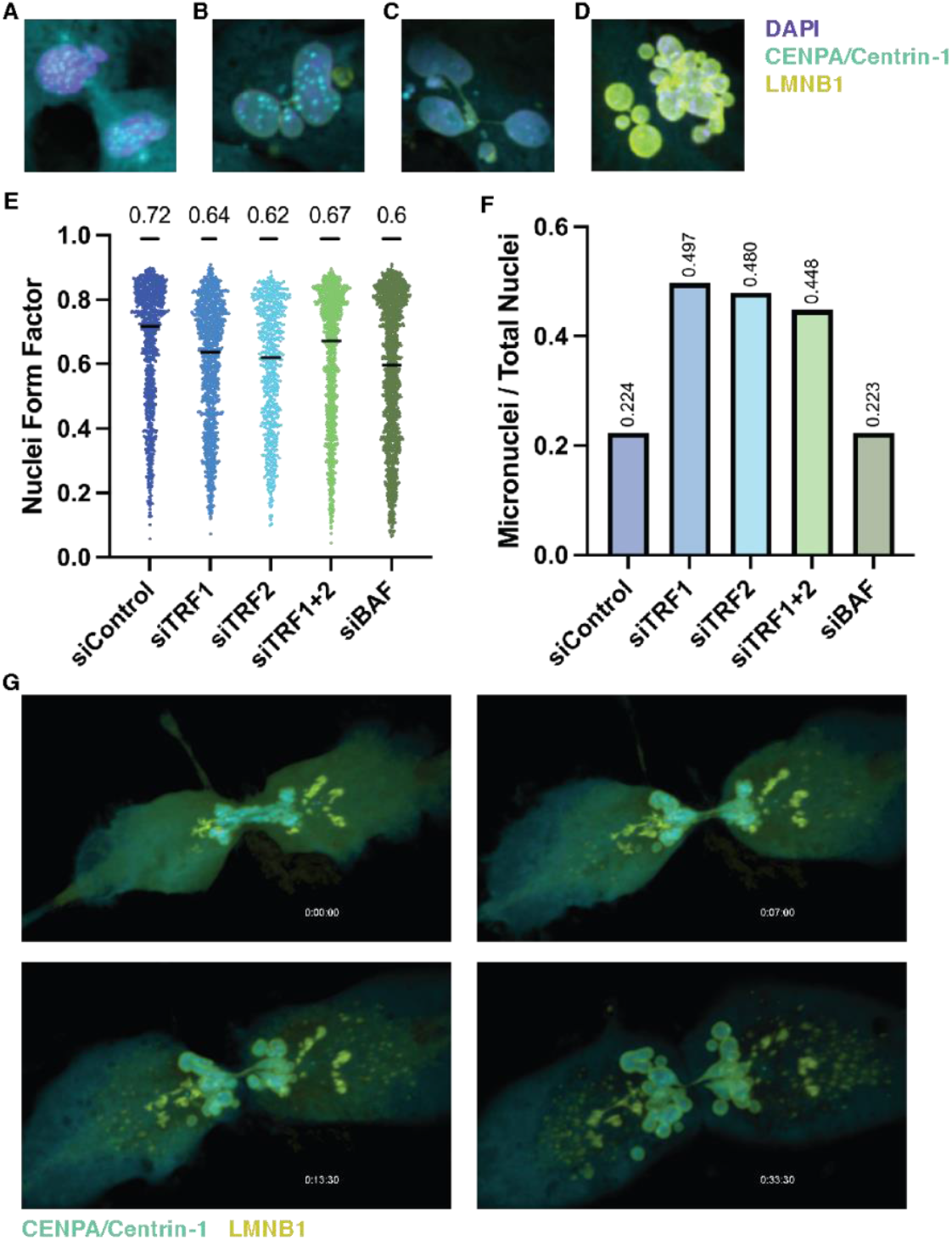
Down-regulation of TRF1 and/or TRF2 results in micronuclei formation. Wide range of phenotypes in cell nuclei is observed after disruption of the shelterin complex through siRNA. DNA in purple, Centromere and Centrioles are labeled in cyan, and Lamin B1 in yellow. Phenotypes ranges from (**A**) bulging of nuclei without separation, (**B**) single micronuclei formation, (**C**) several nuclei and micronuclei with evidence of telomere fusion, to (**D**) severe multinucleation. (**E**) Nuclear form, also known as nuclear circularity, was analyzed in 5 separate treatment groups, each having more than 1000 nuclei. Cells treated with BAF knockdown showed the biggest decrease in nuclear form compared to cells treated with control RNAi. (**F**) Micronuclei was counted using Cell Profiler 3.0, and knockdown of TRF1 and/or TRF2 resulted in the highest instance of micronuclei formation per nuclei. (**G**) Snapshots from movie of RPE-L cell with TRF1 knockdown demonstrating massive amounts of micronuclei formation following mitosis. Centromere and Centrioles are labeled in cyan, and Lamin B1 in yellow.

Randomized nuclear lamina alone would not result in micronuclei formation unless mislocalized protein serves a barrier function for the chromosome it encapsulates. To test the barrier function of our protein, we developed a lagging chromosome to micronuclei assay. First, we treated cells with nocodazole, which destabilizes mitotic spindle and synchronize cells at the G2/M stage. When nocodazole is washed out, synchronized cell proceeds to divide, but has higher instances of lagging chromosome (LC) due to merotelic attachment of microtubules (30). We then performed a systematic assay to test for lagging chromosome versus micronuclei formation. By following induction of LCs during mitosis, we measured distance of each LC and relative LMNB1 levels on the surface of the chromosome (n=13) (Fig 4A). LCs that reached the chromosome mass after LMNB1 accumulation begin were unable to be incorporated into the nucleus, presumably becoming micronuclei (Fig 4B). Like the DNA mass, LCs shows accumulations of LMNB1 and BAF at the telomere tip (movie S10 and S11), however, one outlier was recorded where LMNB1 accumulated before anaphase at a high level with one chromosome, and that chromosome failed to join the chromosome mass despite its close physical proximity to the chromosome mass before LMNB1 accumulated around the DNA mass (Fig 4C, D). Same lagging chromosome to micronuclei assay was done with labeled LAP2B, which surrounds the DNA mass before LMNB1 and is not observed to concentrate on the chromosome tip. No correlation between minimal LAP2B level (occurs before minimal LMNB1 level) and micronuclei formation were observed, however, LC distance being greater than 0 after maximum LAP2B level is reached (occurs after minimum LMNB1 level) resulted in micronuclei formation (n=16), which further validated LMNB1 LC results (Fig S4, movie S12).

**Fig. 4.**
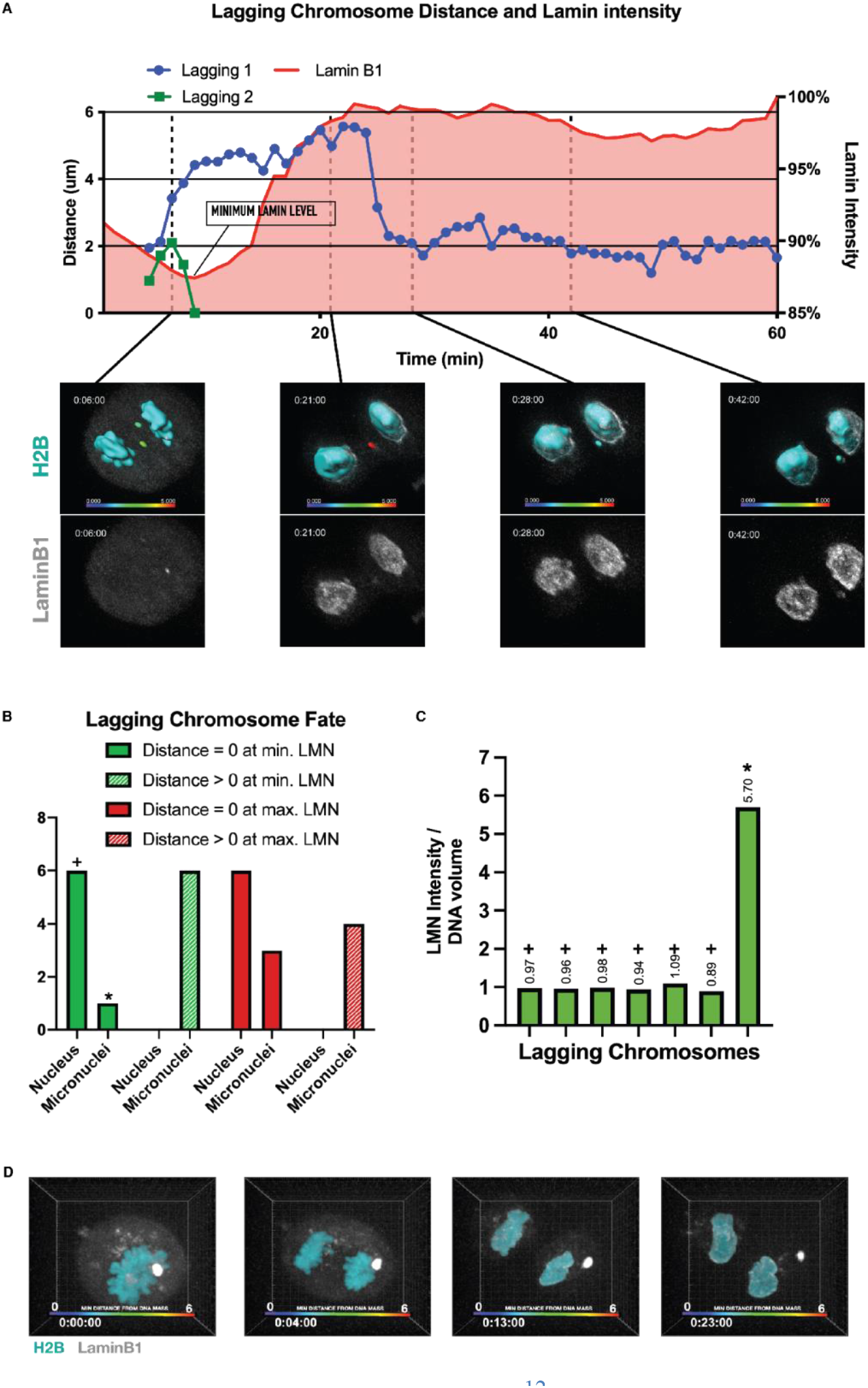
Lamin Accumulation Determines Incorporation of Chromosomes. (**A**) Level of LaminB1 around chromosome mass was quantified, and shown in percentage of maximum lamin level during the live-imaging (shown in red). Distance of lagging chromosome during each time point is calculated by the minimum distance between the lagging chromosome surface and the chromosome mass surface. In example, two lagging chromosomes were present during live imaging, one is represented in blue while the other in green on graph. Images below graph are snapshots from four frames that is represented on the graph in perspective time points. (**B**) Compilation of multiple lagging chromosome from separate movies (n=13). Fate of being incorporated versus forming micronuclei is represented by color vs black. X-axis was set as time relative to the minimum lamin level in each movie. (**C**) Summary of lagging chromosome fate when distance is equal or greater than zero during perspective lamin levels. + shows lagging chromosomes that were incorporated when distance to chromosome mass is 0 while lamin is at minimum level, and * shows one outlier in which the chromosome formed a micronucleus. (**D**) Snapshots of the outlier labelled with * in (**C**). The level of LaminB1 (white) was abnormally and consistently high throughout mitosis.

This study reveals a previously unrecognized role of telomere during the formation of the nucleus serving as an initiation site of NE reassembly during mitosis. Our data show that initial lamin attachment at the telomere region is crucial in ensuring numerous condensed chromosomes are securely packaged into a single nucleus. Telomeres have been shown to have strong correlations with cellular health and aging (31), and our work adds to our understanding of its importance in securing and propagating the full genomic information during the crucial biological process of cell division.

## Supporting information

Materials and Methods, Legends for Figs. S1 to S4, Movie files

U2OS GFP-LaminB1 cells undergoing mitosis,

U2OS GFP-LaminB1 cells undergoing mitosis

U2OS GFP-Sec61B cells undergoing mitosis

taxol and reversine-induced reformation of LMNB1 around the chromosomes to form many micronuclei.

Absence of decondensation in BAF knockdown cells undergoing mitosis

MNB1 accumulates at the telomere tip distal to centromere despite knockdown of BAF

Recombination of telomere in siTRF1-treated cells during cell division

Massive amounts of micronuclei formation following TRF1 knockdown.

BAF concentrating at the chromosome tip in DNA mass and on lagging chromosome in Nocodazole treated cells

Nocodazole treatment induces BAF concentrating at the chromosome tip in DNA mass and on lagging chromosome.

## Acknowledgments

We thank Drs. O. Weiner, S. Dumont, D. Hart, and A. Buchwalter for their comments and suggestions and past and present Mikawa lab members for their suggestions regarding this work. This work is in part of B.Y.

## Author contributions

Conceptualization: PBY, TM

Methodology: PBY

Investigation: PBY, TM

Visualization: PBY

Funding acquisition: TM

Supervision: TM

Writing – original draft: PBY

Writing – review & editing: TM, PBY

## Data and materials availability

All data are available in the main text or the supplementary materials.

## Supplementary Materials

Materials and Methods

Figs. S1 to S3

Movies S1 to S10

Data S1

