## Supplementary material for "Telomere Orchestrates Encapsulation of Mitotic Chromosomes": Materials and Methods, Legends for Figs. S1 to S4, Movie files

**The PDF file includes:**

Materials and Methods
Figs. S1 to S4

**Other Supplementary Materials for this manuscript include the following:**

Movies S1 to S12
Data S1

**Materials and Methods**

Cell Lines / Cell Culture

HeLa cells stably co-expressing Histone H2B-mCherry and Lap2β-EGFP, and HeLA cells stably co-expressing EGFP-BAF and mCherry-Lap2β were generous gift from Dr. Daniel Gerlich (IMBA). RPE1 cells stably expressing CENP-A/centrin1-GFP was a generous gift from Alexey Khodjakov (NY State Department of Health). MCF10AT cell line stably expressing Histone H2B-GFP was a generous gift from Dr. Zev Gartner (UCSF). U2OS GFP-LaminB1 cell line was from Sigma-Aldrich (Sigma-Aldrich, CLL-1033). RPE-L was generated by electroporating mTagRFP-T-LaminB1-10, a gift from Michael Davidson (Addgene plasmid # 58020 ; http://n2t.net/addgene:58020 ; RRID:Addgene_58020), to RPE-18 cells, selected for cells containing plasmid using G418 (Geneticin, Thermo Fisher, 10131035), followed by FACS sorting for desired expression level, and expression of plasmid is maintained by G418 selection every 5 passages.

HeLa and RPE cells were cultured in Dulbecco's modified Eagle medium:Nutrient Mixture F-12 (DMEM:F12; GIBCO) supplemented with 10% fetal bovine serum (FBS; GIBCO), 1% (v/v) penicillin-streptomycin (Sigma-Aldrich), while MCF10AT cells were cultured in Mammary Epithelial Cell Growth Medium Bullet Kit (MEGM; Lonza, CC-3150) at 37°C with 5% CO2 in a humidified incubator.

Immunofluorescence Staining

Cells were grown to 70% confluency, then fixed in 4% paraformaldehyde in PBS for 0.5 hours at room temperature, followed by washing with PBS twice. Blocking media, (1% BSA, 1% triton in PBS) was added for 1 hour at room temperature, followed by primary antibody staining. Primary antibodies were added in blocking solution overnight at room-temperature, followed by 1 hour of PBS washing. Secondary antibodies, were added in blocking solution for 2 hours followed by PBS washing, follow by incubation of PBS with DAPI (1:1000) for 0.5 hours. Following a final wash with PBS, cells were covered with Vectashield mounting media (Vector, H-1000).

Primary antibodies used: Lamin B1 (1:1000) anti-rabbit (Abcam, ab-16048), TRF1 (1:200) anti-mouse (Abcam, ab- 10579), BANF1 (1:200) anti-mouse (Abcam, ab-88464), Lamina B2 (1:50) anti-rabbit (Cell Signaling, D8P3U), and Sec61b (1:200), anti-rabbit (Thermo Scientific, PA3-015).
Secondary antibody used: Alexa Fluor 488 (1:500) anti-rabbit A11008, Alexa Fluor 488 (1:500) anti-mouse A11001, Alexa Fluor 594 (1:500) anti-mouse A21203, Alexa Fluor 594 (1:500) anti-rabbit A11037, Alexa Fluor 647 (1:500) anti-mouse A31571, Alexa Fluor 647 (1:300) anti-rabbit A31573 - all purchased from Thermo Fisher .

Pharmacological Assays

Cells were plated 24hr prior to drug treatment. For lagging chromosome assay, nocodazole (Sigma, M1404) was added to media, and cells were incubated for 16hr. Cells were then washed twice using media, and allowed to proceed pass G2/M phase. For taxol treatment, taxol (Fisher Scientific, NC9019658) was added to media and cells were incubated for 16hr, reversine (Sigma-Aldrich, R3904) was then added to suppress the spindle assembly checkpoint. Final concentration of drugs are as follows: Nocodazole (100 ng/mL) Taxol (585 nM), Reversine (393 nM).

DNA Constructs and siRNA

Cells were electroporated using Thermo Fisher Scientific's Neon Transfection System, following protocol published by Thermo Fisher. Transfection of DNA constructs using lipofectamine were performed using Lipofectamine 3000 (Thermo Fisher), while siRNA were transfected using RNAiMAX (Thermo Fisher), all following protocol by manufacturer.

DNA constructs:

- mTagRFP-T-LaminB1-10 was a gift from Michael Davidson (Addgene plasmid # 58020 ; http://n2t.net/addgene:58020 ; RRID:Addgene_58020)
- mCherry-SEC61B was a gift from Christine Mayr (Addgene plasmid # 121160 ; http://n2t.net/addgene:121160 ; RRID:Addgene_121160)
- Sec61 beta pAcGFP-C1 (802) was a gift from Eric Schirmer (Addgene plasmid # 62008 ; http://n2t.net/addgene:62008 ; RRID:Addgene_62008)
- EGFP-BAF was a gift from Daniel Gerlich (Addgene plasmid # 101772 ; http://n2t.net/addgene:101772 ; RRID:Addgene_101772)
- mCherry-LaminA-C-18 was a gift from Michael Davidson (Addgene plasmid # 55068 ; http://n2t.net/addgene:55068 ; RRID:Addgene_55068)
- Lenti-inducible spCas9, gRNAs for KO TRF1, TRF2, TPP1, POT1, TIN2, and RAP1 were all generous gifts from Dr. Zhou SongYang (Baylor College of Medicine).

siRNA constructs:

- TRF1 siRNA (h) (Santa Cruz Biotechnology, sc-36722).
- TRF2 siRNA (h) (Santa Cruz Biotechnology, sc-38505).
- Ambion siRNA Negative Control #2 (Thermo Fisher Scientific, AM4613).
- Ambion BANF/BAF Silencer Select siRNA (Thermo Fisher Scientific, 4392421).

Microscopy / Image Analysis

For live imaging, cells were plated on glass bottom plates (Ibidi GmbH, 81158), or grow on high precision cover glasses (Deckglaser, 0107052), and allowed to settle for at least 24hr.

Spinning Disk Microscopy were done on a Yokagawa CSU-X1 Spinning Disk Confocal, equipped with a Nikon Perfect Focus system, Sutter Lambda XL lamp, and an Andor Clara interline CCD camera. N-Sim (Structured Illumination Microscopy) were done on a Nikon Ti-E Microscope, equipped with Andor DU897, Nikon Intensilight, and 3D EX V-G 100X/1.49 for SIM gratings. All microscopes were maintained and built by UCSF Microscopy Core, and lenses used were all from Nikon: 100x/1.49 oil APO, and 60x/1.50 oil APO. Image Analysis in 2D and 3D were done on Imaris 9.2 and 9.3 (Bitplane).

Volume Calculation: Optical sections of live images taken with 60x objective were 3D reconstructed. A surface was generated using Imaris 9.3, threshold was kept the same in all movies: 3um for local contrast, 0.3um for surface detail, and bottom 1% of signal (e.g. maximum intensity in movie was 4900, then intensity from 0 to 49) is eliminated while generating the surface. Each frame is 1 minute long, and only the first 30 frames are taken from each movie for quantification to avoid photobleaching affects.

Spread of Lamin Assay: More than 5000 points were randomly generated on red fluorescence areas (lamin was labeled with RFP-t) in each frame, and distance of each point from centromere was calculated. Distance of each dot is plotted on a violin plot, indicating frequency in width of each figure, distance from chromosome tip in length, and measured every 20 second per frame and plotted on x-axis of graph.

Lagging Chromosome to Micronuclei Assay: After incubation and washout of nocodazole (Sigma, M1404), cells were imaged undergoing mitosis using Yokagawa CSU-X1 Spinning Disk Confocal. Surface was generated in Imaris 9.3 using the same method described in volume calculation. Distance between lagging chromosome (LC) and DNA mass was calculated using ShortestDistanceToSurface measurement in Imaris. Intensity level of lamin on surface of lagging chromosome and DNA mass was calculated using intensity of LMNB1 on the surface of DNA mass. LMNB1 intensity and distance between LC:DNA mass were plotted together on the same plot, minimum LMNB1 level and maximum lamin level were then used to categorize the fate of lagging chromosomes.

Nuclear Form and Micronuclei quantification: Cells were transfected with siRNA using method described in DNA Constructs and siRNA. Cells were imaged 48 hours after siRNA treatment. Each treatment group had one large image stitching together 144 images taken with 20x objective. Each image had over 1000 nuclei each, and nuclear form of each nuclei was measured using ImageJ with the FUJI plugin (23), and presence of micronuclei was detected using CellProfiler 3.0 (24).

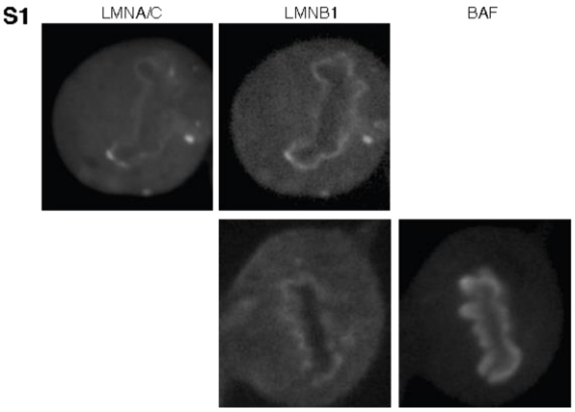


**Fig. S1. Images of cells co-expressing LaminB1 with Lamin A/C or BAF.** Gray scale image showing co-expression of LMNA/C-mCherry with LMNB1-GFP in U2OS cells, and BAF-GFP and LMNB1-RFP-T in RPE-1 cells.

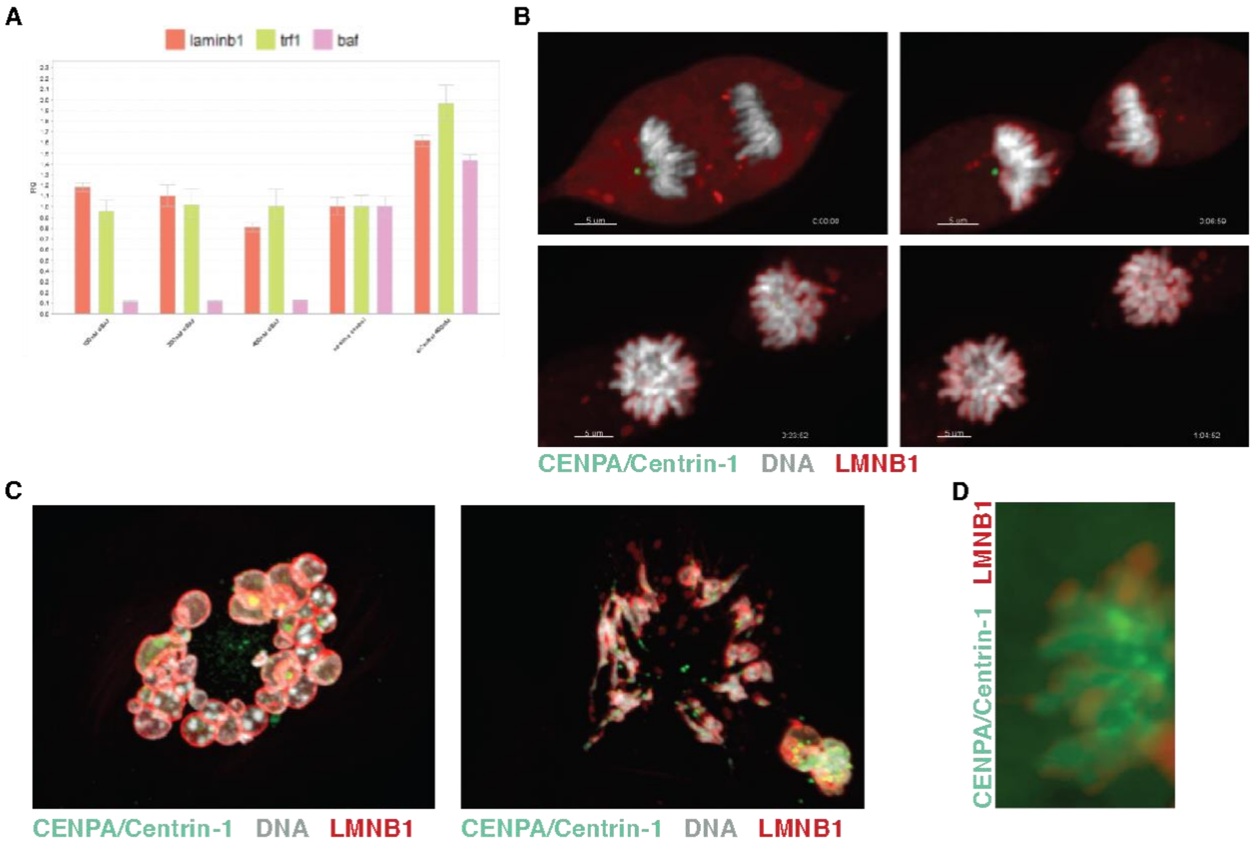


**Fig. S2. BAF Facilitates Chromatin De-Condensation.** (**A**) rtPCR showing effective knockdown of BAF mRNA after 24hr of siBAF. Negative control siRNA showed no significant decrease in BAF mRNA, while 100nM, 200nM and 400nM of siBAF all had significant effect on BAF mRNA level without a decrease in TRF1 or LMNB1 transcription. Transcription levels normalized to GAPDH. (**B**) RPE-L cells 48hr after knockdown of BAF. Normal chromosome (white) de-condensation is not observed in BAF knockdown cells undergoing mitosis. LMNB1 (red) proceeds to spread along each chromosome arm, and normal nuclear morphology is altered following mitosis. (**C**) siBAF inhibited de-condensation of individual chromosomes, and resulted in strand-like micronuclei in cells treated with taxol and reversine. (**D**) siBAF treated RPE-L cell. Snapshot from live imaging of mitosis that shows LMNB1 (red) accumulation at the telomere tip despite knockdown of BAF.

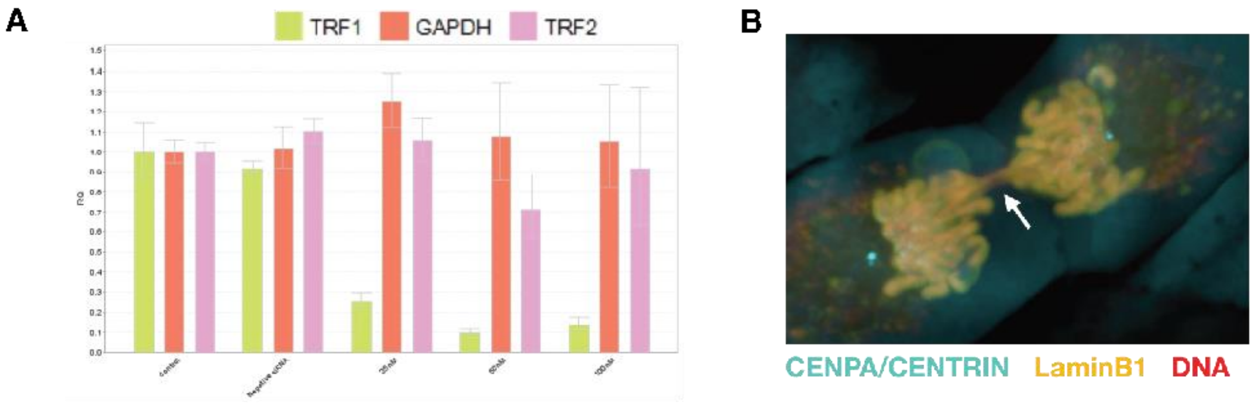


**Fig. S3. Use of siRNA to knockdown Shelterin Complex Unit TRF1.** (**A**) rtPCR showing effective knockdown of TRF1 mRNA after 24hr of siTRF1 transfection. Negative control siRNA showed no significant decrease in TRF1 mRNA, while 25nM, 50nM and 100nM of siTRF1 all had significant effect on TRF1 mRNA level, while TRF2 and GAPDH remained normal. Transcription levels normalized to Beta-Actin. (**B**) After 48hr of siTRF1, recombination of telomere can be observed in dividing cells, indicating delocalization of shelterin complex and structural change of telomere D-loop.

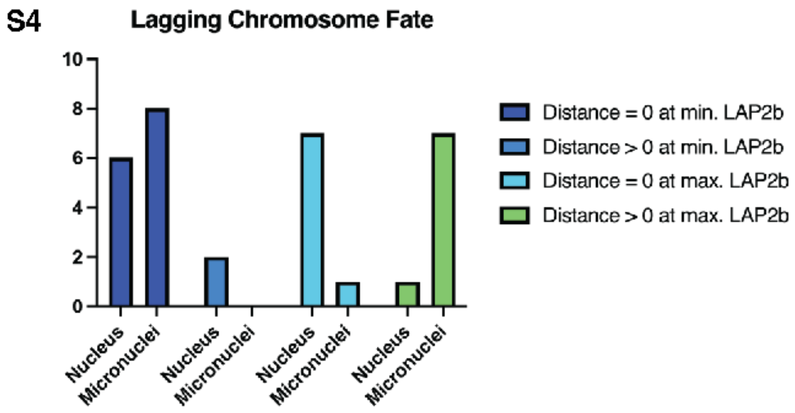


**Fig. S4. Use of siRNA to knockdown Shelterin Complex Unit TRF1.** Summary of lagging chromosome fate when distance is equal or greater than zero during perspective Lap2β levels. Minimum LAP2B level and Maximum LAP2B level groups are describing the same LC (n=16).

**Movie S1.**

U2OS GFP-LaminB1 (Cyan) cells undergoing mitosis, DNA labeled with SiR-DNA (Red). Example of volume calculation using IMARIS is shown as a surface is generated based on the SiR-DNA labeling (Red channel).

**Movie S2.**

U2OS GFP-LaminB1 (Cyan) cells undergoing mitosis, DNA labeled with SiR-DNA (Red). Movie is paused during late anaphase to show Lamin concentrating at the chromosome tip.

**Movie S3.**

U2OS GFP-Sec61B (Cyan) cells undergoing mitosis, DNA labeled with SiR-DNA (Red).

**Movie S4.**

HeLA EGFP-BAF (Cyan) mCherry-Lap2βeta (Yellow) cells undergoing mitosis, DNA labeled with SiR-DNA (Red). Movie is paused during early telophase to show BAF concentrating at the chromosome tip.

**Movie S5.**

RPE-L, CENP-A/centrin1-GFP (Green) and LMNB1-RFP-T (Red) were treated with taxol and reversine and live-imaging shows reformation of LMNB1 around the chromosomes to form many micronuclei.

**Movie S6.**

RPE-L cells 48hr after knockdown of BAF. Normal chromosome (white) de-condensation is not observed in BAF knockdown cells undergoing mitosis. LMNB1 (red) proceeds to spread along each chromosome arm, and normal nuclear morphology is altered following mitosis.

**Movie S7.**

siBAF treated RPE-L cell undergoing mitosis. LMNB1 (red) accumulates at the telomere tip distal to centromere (Green) despite knockdown of BAF.

**Movie S8.**

After 48hr of siTRF1, recombination of telomere can be observed in dividing cells, indicating delocalization of shelterin complex and structural change of telomere D-loop.

**Movie S9.**

RPE-L cell with TRF1 knockdown demonstrating massive amounts of micronuclei formation following mitosis. Centromere and Centrioles are labeled in cyan, and Lamin B1 in yellow.

**Movie S10.**

HeLA EGFP-BAF (Cyan) mCherry-Lap2βeta (Yellow) cells treated with nocodazole and undergoing mitosis after wash, DNA labeled with SiR-DNA (Red). Lagging chromosome was induced, and movie is paused during early telophase to show BAF concentrating at the chromosome tip in DNA mass and on lagging chromosome.

**Movie S11.**
MCF10AT (Gray) with exogenous expression of LMNB1-RFP-T (Red) was treated with nocodazole and undergoing mitosis after wash. Lagging chromosome was induced, and movie is paused during early telophase to show BAF concentrating at the chromosome tip in DNA mass and on lagging chromosome.

**Movie S12.**
HeLa cells stably co-expressing Histone H2B-mCherry (Yellow) and Lap2β-EGFP (Cyan) was treated with nocodazole and undergoing mitosis after wash. Lagging chromosome was induced, and movie is paused to show the overlying of surface to DNA mass and Lap2β-EGFP using IMARIS for analysis.

**Data S1.**
Images of the 5 treatment groups used for analysis in FIJI and Cell Profiler 3.0.
